# A Neurofeedback therapy of facial expression recognition in Autism shifts connectivity to higher levels within the third visual pathway in relation to clinical improvements

**DOI:** 10.64898/2026.04.14.718381

**Authors:** Bruno Direito, Alexandre Sayal, Susana Mouga, Miguel Castelo-Branco

## Abstract

The mechanistic role of the third visual pathway in autism spectrum disorder (ASD) remains unknown. We previously developed a neurofeedback therapy for autism targeting the posterior superior temporal sulcus (pSTS), a region in this network that underlies the perception and imagery of emotional facial expressions, resulting in improvements in adaptive behavior and recognition of fear in facial expressions. Here, we investigated the impact of this 5-session therapy on the functional connectivity of that core region of the third visual pathway. We found evidence for a profound reorganization of this network with treatment-induced decreases in connectivity between low-level visual areas, the pSTS, and the posterior occipital face area (OFA), and increased connectivity with higher-level visual regions (fusiform face area - FFA), right middle STS (mSTS), and parietal cortex. These changes, suggesting the restoration of connectivity in regions known to be underconnected in ASD, such as mSTS and pSTS, and in a set of regions belonging to the temporoparietal junction and the ventral attention network, which are known to be involved in broader aspects of social cognition, were positively associated with clinical improvements. The demonstration of treatment response associated with network reconfiguration paves the way for multicentric trials to probe this observed reorganization as a treatment target.

## Introduction

### Background

Autism spectrum disorder (ASD) is a life-long neurodevelopmental condition characterized by difficulties in social communication and interaction, as well as restricted/repetitive patterns of behavior, interests, or activities, as outlined in the DSM-5-TR (American Psychiatric Association, 2022). Despite these core features, ASD is characterized by significant etiological and phenotypic heterogeneity(Mouga et al., 2016), reflecting the complex genetic, environmental, and neurobiological factors underlying the condition. Understanding how activity and connectivity patterns vary is crucial for developing a better grasp of both diagnostic and therapeutic outcome biomarkers.

In terms of imaging markers, ASD has been linked to atypical function in core social brain regions and altered connectivity patterns (Just et al., 2012). In particular, reduced attention to expressive facial features is a frequently replicated finding in research involving individuals with ASD (Harms et al., 2010), with direct implications for understanding the brain networks underlying social cognition.

The seminal earlier work of (Haxby et al., 2000) identified the fundamental functional architecture of face-processing networks: the fusiform gyrus (FG) is more involved in processing invariant facial features, such as identity, whereas variable aspects, such as eye gaze and expression, recruit the posterior superior temporal sulcus (pSTS). (Fox et al., 2009) further mapped a set of core and extended face-selective regions involved in face processing. The fusiform face area (FFA), located in the right fusiform gyrus, the inferior occipital gyrus (known as the occipital face area, OFA), and the posterior superior temporal sulcus (pSTS), are currently regarded as the “core system” for face processing. Beyond the core system, several additional regions contribute to face perception, forming the “extended system.” These regions include the inferior frontal gyrus (IFG), the amygdala (AMG), the precuneus (preC), the anterior paracingulate (aPC) gyrus, and a more anterior portion of the STS (aSTS). Ultimately, (Fox et al., 2009) proposed a network comprising six regions in the core system and eight regions in the extended system.

Among the functional neural alterations identified in ASD regarding either facial identity processing or facial expression recognition, the latter stands out, with atypical cortical hypoactivation patterns across various tasks. These findings highlight key changes in regions such as the FG and the pSTS, which are central in processing facial expressions and social cues (Shih et al., 2011).

Recently, a third visual pathway specialized in social perception has been proposed, emphasizing the critical role of the right STS in processing socially relevant cues and supporting high-level social perception (McMahon et al., 2023; Pitcher & Ungerleider, 2021). This pathway complements the classical ventral (what) and dorsal (where/how) visual streams in a third visual stream on the lateral surface of the brain, specialized for processing social information. This third stream also exhibits hierarchical organization and is involved in higher-level aspects of interpreting dynamic social stimuli, social interactions, and affective content. In this context, the posterior and middle regions of the STS (pSTS and mSTS) are considered pivotal hubs for integrating visual and social information. Preliminary findings by (Li et al., 2025) suggest a disruption of this social visual pathway in ASD, with significant underconnectivity observed between the pSTS and mSTS in the right hemisphere in children with ASD compared to typically developing children. These results highlight the importance of investigating the functional integrity of this pathway to better understand the neural mechanisms underlying social deficits in ASD and to inform targeted intervention strategies.

Neuroimaging-based therapeutic interventions, such as neurofeedback (NF), have been advocated as a novel approach to target atypical activation patterns by modulating the underlying brain networks. The promise of NF in revealing principles of cognition while targeting abnormal local and global network patterns rests on its approach to causal modulation. In this sense, by providing individuals with real-time feedback on their brain activity, NF allows researchers to target neural processes through a region-specific node. In particular, these interventions can aim to restore typical functional connectivity and improve social cognition in individuals with ASD. In (Pereira et al., 2024), we have shown differences in connectivity associated with NF training of executive function in ASD, with potential impacts on whole-brain activity levels. Network neuroscience aims to model brain networks using the principles and tools of graph theory, enabling the quantitative characterization of networks and their nodes. In this sense, NF targeting the behavior of a specific node has the potential to affect the entire network, and these tools allow us to characterize this interaction (Bassett & Khambhati, 2017).

In the present study, we investigated changes in functional connectivity (FC) following a multi-session, real-time fMRI neurofeedback (NF) clinical trial. The NF treatment targeted the pSTS, a key area of the third visual pathway, and we specifically investigated whether FC modifications occurred in relation to this core region.

In this context, a major research question is whether NF interventions induce adaptive FC changes in ASD. This study investigates FC changes observed in participants from clinical trial NCT02440451, which reported beneficial outcomes such as improved discrimination of fear recognition in emotional faces (Direito et al., 2021).

Specifically, we investigated task-based functional connectivity considering the individual functional localizer of the NF target pSTS, and the NF imagery task between the last and first runs of treatment sessions, addressing two primary research questions: First, does the treatment induce changes in ROI-to-ROI connectivity within the extended face network, as defined in (Fox et al., 2009). Second, and most importantly, does the pSTS, a core region of the third visual pathway and our NF target, shift its connectivity within this pathway? We hypothesize that NF training may influence the FC patterns of this network.

In testing this hypothesis, we aim to characterize changes associated with the modulation of a specific target neurofeedback region involved in facial recognition and expression. This analysis seeks to determine the brain regions that present altered connectivity before vs. after NF, which are directly linked to the target pSTS region.

Importantly, we further investigated the association between FC changes and clinical features, particularly those associated with social adaptive behavior. We hypothesized that FC after training would change within the third visual pathway and that connectivity strength would be associated with the severity of autism symptoms.

## 2. Methods

### 2.1. Subjects and experimental procedure

Fifteen male participants (19 years and 11 months, ±3 years and 3 months) with ASD enrolled in a 5-session training program of real-time fMRI NF targeting the processing of emotions in facial expressions, using the pSTS as the target ROI, in a previously published trial NCT02440451 (Direito et al., 2021).

The primary outcome measure for the intervention was the Facial Expressions of Emotion—Stimuli and Tests—FEEST (The Emotion Hexagon test) (Young et al., 2002). The Emotion Hexagon test involves recognition of six basic emotions: anger, disgust, fear, happiness, sadness, and surprise. The test was taken at the first visit and the last imaging session. The patients were also characterized before and after the NF training with the Autism Treatment Evaluation Checklist (ATEC) (Rimland & Edelson, 1999). The ATEC is a caregiver-administered questionnaire designed to assess changes in ASD severity in response to treatment. Caregivers were blind to the fact that this instrument was used as a secondary outcome measure.

Each session included a functional localizer to create a mask for the pSTS, defining the target area, followed by four imagery task-based runs - a training run without feedback, two NF runs, and a transfer run (also without feedback). During the NF, training, and transfer runs, participants were instructed to imagine facial expressions corresponding to happy, sad, and neutral emotions. Feedback was provided via a visual stimulus that morphed an avatar’s face in response to modulation of pSTS activity (Direito et al., 2019).

The functional localizer was performed to identify the target region of interest (ROI) using a block-based design consisting of 40 blocks of 8 seconds each, incorporating five different conditions: Neutral - static neutral faces to control for static aspects of face processing; Moving dots - randomly moving dots to control for motion processing in the pSTS; and Happy, Sad, Alternate expressions - morphing facial expressions. The primary contrast of interest involved the balanced subtraction of the Neutral and Moving dots conditions from the emotion expression conditions (Happy, Sad, and Alternate). Each imagery run consisted of 25 blocks (12 regulation blocks featuring imagery of non-neutral expressions alternating with 13 regulation blocks of neutral expression imagery), each block with a duration of 24 seconds (see Supplementary Methods for further details).

The target ROIs of each participant were defined during the MRI acquisition: using real-time processing software, a three-dimensional box was manually placed over the right pSTS voxels that showed the strongest response in the statistical activation map contrasting Happy, Sad, and Alternate > Neutral and Moving dots conditions. The anatomical definition of the first sulcus, inferior to the lateral fissure, as described by (Deen et al., 2015), guided the selection. This procedure was performed across all five sessions, ensuring precise, subject-specific localization in each.

This dataset has been obtained in a therapeutic trial investigating clinical and behavioral effects as well as the modulation of the target region during NF training (Direito et al., 2021)-

### 2.2. Data acquisition

Data was acquired on a 3T Siemens Magnetom Tim Trio scanner with a 12-channel head coil, at the Portuguese Brain Imaging Network. Each scanning session started with a T1-weighted high-resolution magnetization-prepared rapid acquisition gradient echo sequence for co-registration of functional data (176 slices; TE: 3.42 ms; TR: 2530 ms; voxel size 1 mm^3^ isotropic, FA: 7°; matrix size: 256 × 256). The functional data were acquired using an echo planar imaging sequence (160 volumes, TR = 2000 ms, TE = 30 ms, flip angle = 75°, 32 slices, matrix size 64 × 70, in-plane voxel size = 3 × 3 mm^2^, slice thickness = 2.5 mm, slice gap = 0.5 mm).

### 2.3. Data preprocessing

Data were preprocessed and analyzed using the CONN release 22.a (Nieto-Castanon, 2020) and SPM release 12.12.6. The preprocessing pipeline defined in these tools comprised functional realignment using realign & unwarp, slice-timing correction, outlier identification using ART as acquisitions with framewise displacement above 0.9 mm or global BOLD signal changes above five standard deviations, normalization into MNI space, segmentation into gray matter, white matter, and CSF and spatial smoothing with a Gaussian kernel of 8 mm full-width half maximum (FWHM). Physiological artifacts and residual subject movement effects were removed through a combination of linear regression of potential confounding effects in the BOLD signal (noise components from cerebral white matter and cerebrospinal areas, estimated subject-motion parameters, identified outlier scans, and task effects) and temporal high-pass filtering (0.008 Hz cutoff). For additional details, please refer to Supplementary Materials.

### 2.4. Functional connectivity analyses

Our primary goal is to characterize differences in connectivity associated with NF training by comparing connectivity patterns at baseline (first session) and after the training sessions (last session). To this end, we analyze the data from the localizer runs - a protocol designed to map the dynamical facial expression perception network - and the transfer runs (imagery task without feedback, performed to evaluate the impact of feedback). We followed two approaches: connectivity analysis within the extended face network and seed-to-voxel analysis using the target region as the seed.

#### 2.4.1. ROI-to-ROI

The extended network for face processing was defined based on (Fox et al., 2009) and consists of 14 areas (6 for the core system and 8 for the extended): the right and left FFA (located in the fusiform gyrus), the right and left OFA (inferior occipital gyrus), and right and left pSTS; right and left IFG, right and left AMG, the preC, the aPC gyrus, and right and left mSTS. Each ROI was defined as an 8 mm radius sphere centered in the average coordinates of each area (Talairach to MNI conversion was performed on the Bioimage Suite (Lacadie et al., 2008)) (Table 1).

**Table 1.** MNI coordinates of the extended face processing network.

| **Region-of-interest (extended face network)** | **x** | **y** | **z** |
| --- | --- | --- | --- |
| Right OFA | 35 | -79 | -22 |
| Left OFA | -38 | -74 | -25 |
| Right FFA | 38 | -47 | -29 |
| Left FFA | -40 | -42 | -28 |
| Right pSTS | 56 | -41 | -1 |
| Left pSTS | -54 | -52 | 2 |
| Right mSTS | 52 | -2 | -18 |
| Left mSTS | -58 | -14 | -12 |
| Right AMG | 19 | -2 | -17 |
| Left AMG | -19 | -4 | -21 |
| Right IFG | 48 | 21 | 21 |
| Left IFG | -49 | 18 | 21 |
| preC | 1 | -65 | 31 |
| aPC | 7 | 57 | 23 |

The 2nd-level, ROI-to-ROI, analyses with the 14 ROIs consisted of a between-sessions contrast (sessions 5 > session 1), paired t-test, considering the facial expressions morphing conditions (p < 0.05, uncorrected at connection-level, with a p < 0.05 false discovery rate (FDR) correction at network-level (Zalesky et al., 2010)). This test allowed us to detect task-related connectivity changes within the extended face processing network from session 1 to session 5.

We also performed an ROI-to-ROI analysis on the transfer runs (comparing session 1 and session 5) using the same network and second-level contrast, focusing on the conditions involving facial expression imagery.

To capture changes in the graph network supporting the data, we computed the Clustering coefficient using the correlation coefficient as the network edge (threshold of 0.15), a measure of local integration that characterizes each node as the proportion of connected edges in its local neighborhood.

#### 2.4.2. Seed-to-Voxel

The seed region for the seed-to-voxel analysis was selected per participant based on a fixed-effects analysis of the localizer runs, which were designed to define the pSTS, performed throughout the neurofeedback training sessions. We used the contrast “facial expressions morphing > static face + moving dots” to identify the clusters associated with dynamic face processing and selected the one on the posterior portion of STS (Direito et al., 2019, 2021; Simões et al., 2020). To standardize the seed shape and size for each participant, we created a spherical mask with a radius of 8 mm centered on the peak voxel of the cluster for each participant.

The Seed-to-Voxel 2nd-level analyses consisted of a between-sessions contrast (end versus beginning of treatment; sessions 5 > session 1), paired t-test, considering the facial expressions morphing conditions (p < 0.05, uncorrected at connection-level, with a p < 0.05 false discovery rate (FDR) correction at cluster-size). In other words, we asked which voxels show a connectivity change to the seed region, from the very beginning to the end of treatment, during facial expression morphing.

#### 2.4.3. Relation between functional connectivity and behavioral outcomes

To investigate the association between intervention-induced changes in functional connectivity and clinical outcomes, we correlated connectivity changes with the corresponding changes in behavioral ATEC scores.

## 3. Results

### 3.1. Connectivity analysis of neurofeedback training impact on the face network

#### 3.1.1. ROI-to-ROI analysis

Significant decreases in connectivity were found between the bilateral OFAs and the right pSTS, and the left OFA and left pSTS (Figure 1). The results also show increased connectivity between the right mSTS and two areas: the left pSTS and the left FFA. Overall, these results suggest a shift from low to higher-level connectivity within the third visual pathway.

**Figure 1.**
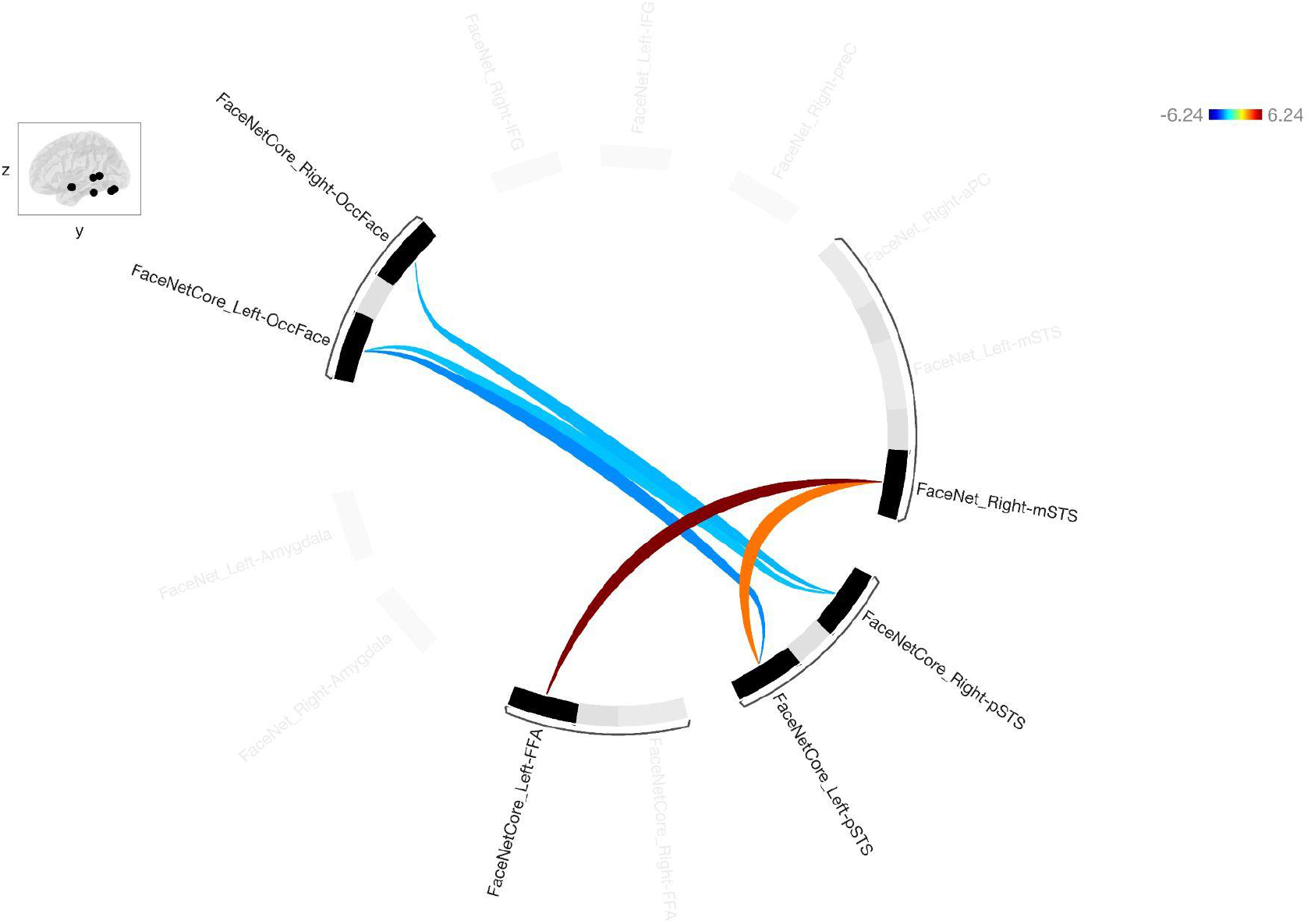
Group ROI-to-ROI connectivity analysis considering the localizer run, comparing last > first neurofeedback session, involving the extended face network (p < 0.05, uncorrected at connection-level, with a p < 0.05 false discovery rate (FDR) correction at network-level (Zalesky et al., 2010)). The edges connecting pSTS and low-level face processing regions (OFA) decrease, while higher-level connectivity with FFA and connectivity between right mSTS and left STS increase.

Network analysis results show an increase in clustering coefficient (characterizing the proportion of connected edges within a node’s neighboring sub-graph) of the right pSTS region (p=0.019, uncorrected, because the pSTS is the focus of our hypothesis).

#### 3.1.2. Seed-to-Voxel analysis and parametric changes of connectivity over time

Comparing the connectivity at the end of the last treatment session and the beginning of the first treatment session session, 2nd level analysis showed significant reductions in FC from the seed (pSTS target region for each participant) to bilateral Lateral Occipital Cortex and Occipital Fusiform Gyrus, in line with the ROI-to-ROI analysis, PreCentral gyrus (right), Frontal Pole (left) and increased FC to the Parahippocampal Gyrus (anterior portion, Left) (Figure 2).

**Figure 2.**
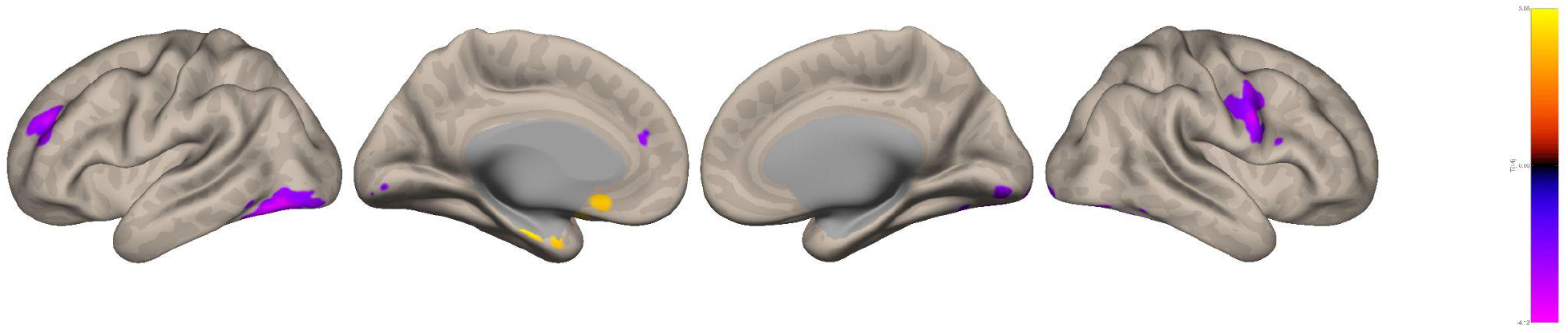
Group seed-based (considering the pSTS as the seed region) connectivity analysis, comparing last > first neurofeedback session (p < 0.05, uncorrected at connection-level, with a p < 0.05 false discovery rate (FDR) correction at cluster-size), highlighting decreased connectivity with low-level visual areas and increased connectivity with the Parahippocampal Gyrus.

Finally, we identified changes across the training sessions. To this end, we used data from the localizer run across the five sessions and explored patterns that showed parametric variation over time. The results show a decrease in connectivity over time between the target region and the bilateral Occipital Fusiform Gyrus, Lateral Occipital Cortex, and Inferior Temporal Gyrus (Figure 3).

**Figure 3.**
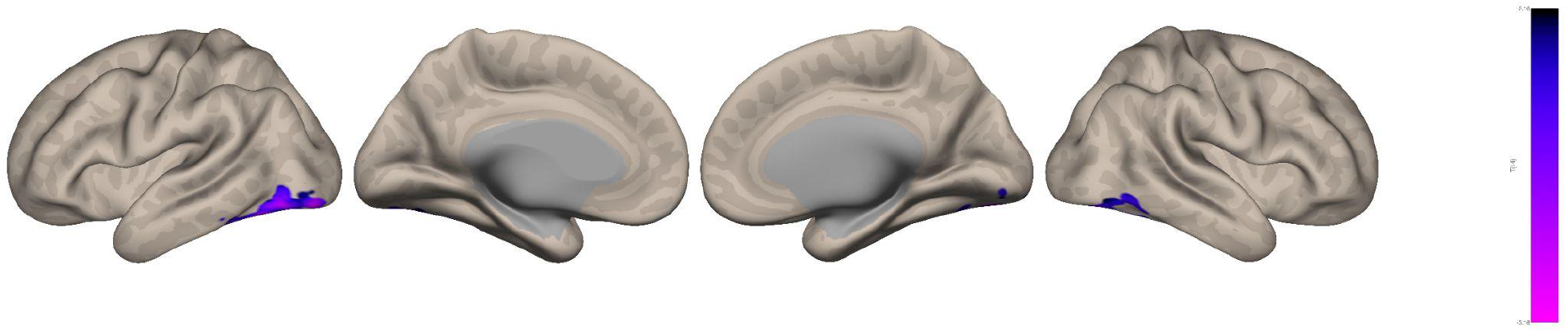
Group seed-based (considering the pSTS as the seed region) connectivity analysis, displaying a regression considering the five sessions (p < 0.05, uncorrected at connection-level, with a p < 0.05 false discovery rate (FDR) correction at cluster-size) and highlighting a decreased connectivity over time between the pSTS and Occipital Fusiform and inferotemporal areas.

#### 3.1.3. Relation between functional connectivity and clinical outcomes

When correlating improvements (i.e., *a decrease*) in ATEC scores with the FC change (between sessions 5 and 1), we observed a significant negative correlation between the target pSTS to right Supramarginal Gyrus - posterior division, Superior Temporal Sulcus, Middle Temporal Gyrus - temporo occipital part, Angular Gyrus, and other regions of the Temporo-Parietal Junction. All these regions belong to the social cognition and theory of mind networks. We found a positive correlation between improvements in ATEC scores with increased FC from pSTS to PreC, PC (Cingulate Gyrus, posterior division), MTG (Left anterior and posterior divisions), left Temporal Pole, and Paracingulate Gyrus (Figure 4). Intriguingly, these regions largely relate to the default mode network.

**Figure 4.**
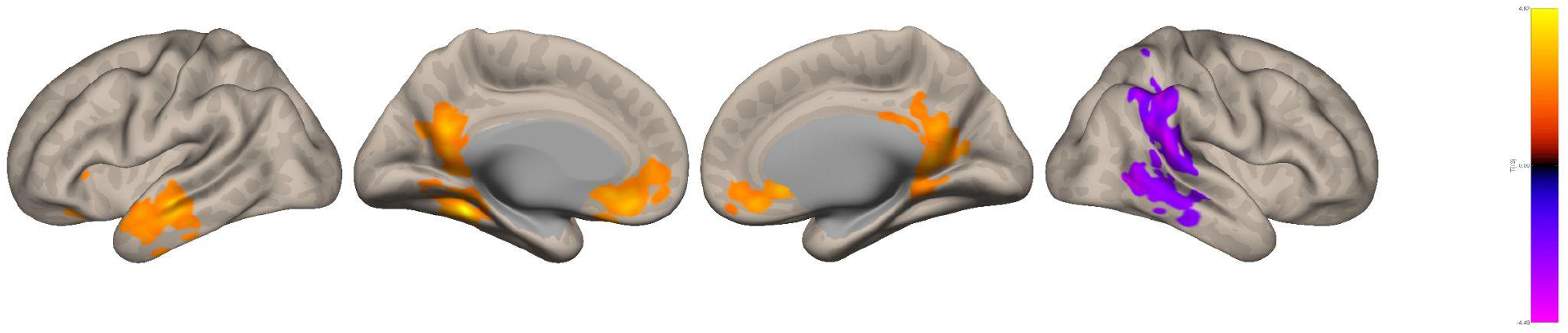
Improvement related increases in the social cognition network with decoupling from the default mode network (DMN) Group seed-based (considering the pSTS as the seed region) connectivity analysis, displaying correlation between changes in connectivity (last > first neurofeedback session) with the corresponding change in ATEC scores (lower scores meaning improvement) (p < 0.05, uncorrected at connection-level, with a p < 0.05 false discovery rate (FDR) correction at cluster-size).

### 3.2. Functional connectivity changes during imagery after neurofeedback training

#### 3.2.1. Seed-to-Voxel

We tested for differences in task-based functional connectivity associated with the imagery of facial expression morphing during the final transfer run between the first and last sessions. The results show a decrease in connectivity between the seed region (target of neurofeedback intervention) and the bilateral Frontal Pole, left Lateral Occipital cortex, left Supramarginal gyrus, and left angular gyrus, bilateral Parahippocampal gyrus, and left Middle Frontal Gyrus (Figure 5).

**Figure 5.**
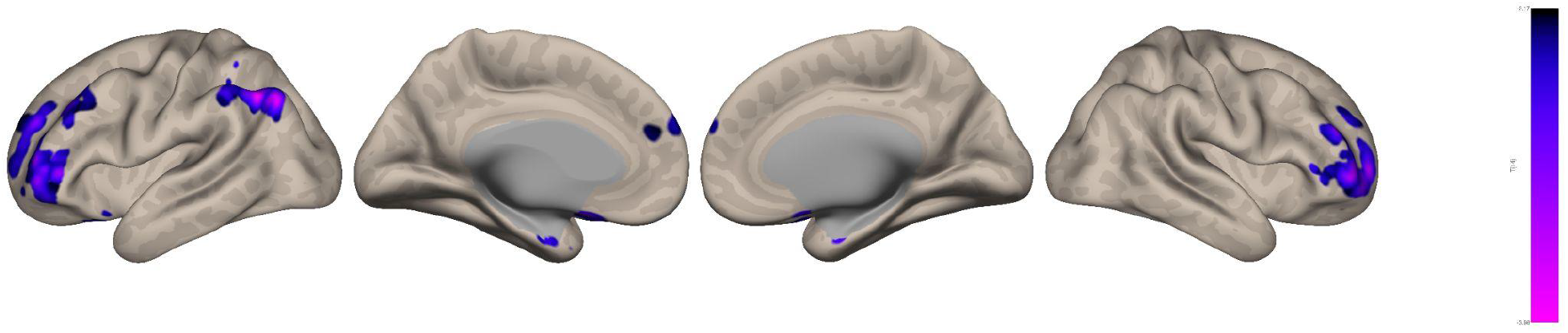
Group seed-based (considering the pSTS as the seed region) connectivity analysis considering the last transfer run, comparing last > first neurofeedback session (p < 0.05, uncorrected at connection-level, with a p < 0.05 false discovery rate (FDR) correction at cluster-size), highlighting decreased connectivity with frontal areas.

#### 3.2.2. Relation between functional connectivity and behavioral outcomes

When correlating improvements in ATEC scores with the functional connectivity change (between sessions 5 and 1), we observed a significant increase with the right Lingual Gyrus, bilateral Occipital Pole, bilateral Lateral Occipital Cortex, Precuneous, bilateral Occipital Fusiform Gyrus, left Temporal Occipital Fusiform Cortex, posterior Cingulate Gyrus, bilateral temporo-occipital junction, right Temporal Occipital Fusiform Cortex, right posterior Supramarginal Gyrus, left Thalamus, bilateral Hippocampus and right Parahippocampal gyrus (Figure 6). In sum, improvements in ATEC were associated with changes in the connectivity of visual, social cognition, and DMN networks.

**Figure 6.**
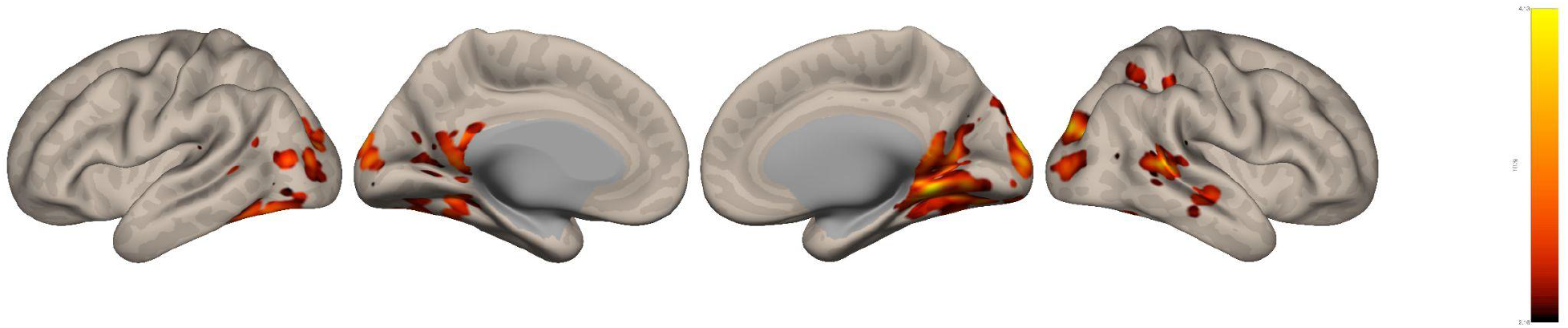
Group seed-based (considering the pSTS as the seed region) connectivity analysis considering the transfer run, displaying correlation between changes in connectivity (last > first neurofeedback session) with the corresponding change in ATEC scores (p < 0.05, uncorrected at connection-level, with a p < 0.05 false discovery rate (FDR) correction at cluster-size).

## 4. Discussion

Here, we used longitudinal data from a 5-session clinical trial to measure functional connectivity before and after neurofeedback training of the pSTS, using task-based fMRI. We tested the hypothesis that regions connected to this target face expression processing area would change their connectivity in a treatment response-associated manner. We found that connectivity patterns in the face-processing network were associated with patients’ clinical outcomes. Specifically, our findings indicate that neurofeedback training may have led to variations in connectivity patterns associated with the pSTS, a key area within the third visual pathway.

We initially assessed pre- to post-training differences in the connectivity of the face network using data from the functional localizer, designed to map the neural correlates of dynamic facial expressions (Direito et al., 2019). We hypothesized associations between changes in clinical scores (ATEC) and the extended facial expressions brain network (Fox et al., 2009). We also used an imagery task, similar to the neurofeedback training task but with the feedback interface removed, to examine connectivity differences between the beginning and end of the NF training protocol.

Our findings suggest that NF training induces changes in the connectivity profile of the extended face network. Importantly, these changes are not limited to the target region but involve the functional network associated with the processing of facial expressions.

### Changes in connectivity associated with NF training

Understanding the neural mechanisms underlying the effects of the NF training process and their association with neurobehavioral outcomes was the main goal of this study. Research indicates that neurofeedback involves changes in the interaction across brain networks, including those associated with self-regulation, the specific cognitive task being targeted, and feedback learning processes (Sitaram et al., 2016; Skottnik et al., 2019). Significant progress in measuring interactions across diverse organizational scales - from molecular to neural populations and major brain networks - has revealed that rewiring of network’ edges underpins changes in behavioral patterns (Bassett & Sporns, 2017). Most studies monitor modulation in the target region(s) or target connection(s), but from a network perspective, it is important to understand how modulation at a network node contributes to the function of other network elements in health and disease (Bassett & Khambhati, 2017).

### NF training using pSTS as the target and the impact on the extended face network

We found decreased connectivity between the bilateral OFAs and the right pSTS, and the left OFA and left pSTS. This suggests decreased connectivity within low-level nodes of the face-processing network. Considering the impairment in facial expression recognition in ASD, previous studies have established that structural and functional connectivity is altered in this network, including pSTS, FG, orbitofrontal cortex, and amygdala (Alaerts et al., 2014). The pSTS also provides strong visual input to fronto-parietal regions implicated in ASD (Simões et al., 2020). Altogether, these results suggest poor integration among areas serving visual, attentional, and socio-emotional functions (Williams et al., 2006). The decrease in the FC between pSTS and the OFA, may imply a dissociation between the first stage in the hierarchy of the face perception network and the integration of facial emotion expression signals in the pSTS. This decrease is combined with an increase in connectivity between the right mSTS and two areas: left pSTS and left FFA. (Li et al., 2025) investigated the notion of a social visual pathway, reporting a decrease in connectivity between right pSTS and mSTS in children with ASD. Our results suggest that NF training may contribute to an increase in this functional connection, reinforcing processing within the third, social information processing, and visual pathway. It is remarkable that positive changes in connectivity of regions known to be underconnected in ASD, such as mSTS and pSTS, and a set of regions belonging to the temporoparietal junction underlying broader aspects of social cognition, were associated with clinical improvements.

The results of the seed-to-voxel analysis, considering the NF training runs, support the notion that there is a dissociation between pSTS as the emotion integration hub and processing in earlier visual areas associated to face processing; and a reinforcement of the functional connection between pSTS and the anterior portion of the Parahippocampal Gyrus (Zeidman & Maguire, 2016), possibly linked to more high-level emotion processing and visual recognition memory networks.

The parametric trend analysis, considering the 5 sessions, showed decreased connectivity between the seed and the ventral, most posterior parts of the inferotemporal pathway. Altogether, our data may suggest that NF training reinforces the “social stream”, reinforcing pSTS-mSTS connections and dissociating from the earlier parts of the visual, “what” stream.

### FC-Behavioral analysis

To investigate the potential relation between the clinical changes and FC, we performed a correlation analysis between the variation of the FC between the first and last sessions of treatment and the improvement, meaning a decrease, in the ATEC scores (before NF-training vs. after). The results show a negative correlation between ATEC decreases (meaning clinical improvement) and connectivity between pSTS and nearby structures, the inferior parietal lobule (IPL), with a right lateralization, suggesting a treatment-related increase in parietal connectivity. On the other hand, ATEC decrease presents a positive correlation with the FC between the seed (right pSTS) and Precuneus PC (Cingulate Gyrus, posterior division), and left MTG (anterior and posterior divisions), left temporal pole, and Paracingulate Gyrus, suggesting a treatment-induced decoupling from the default mode network. These data suggest a reinforcement of the social pathway, particularly the pSTS-mSTS connection.

Impairments in connectivity between pSTS and IPL and their connections to frontal regions have been previously shown in task-based fMRI analysis in ASD (Alaerts et al., 2014) and represent the backbone of the ToM theory on the pathophysiology of ASD - pSTS hypoconnectivity to fronto-parietal regions supports this hypothesis. These results suggest that NF training may have supported a functional change, reinforcing the social pathway and frontal projections.

The hyperactivation of the Precuneus and parietal regions has been linked to increased attentional load (Simoes et al., 2018; Wang et al., 2004), and increased connectivity may represent an effort from the network to adjust. The activation/modulation of the activity in the precuneus has also been shown in different NF training studies involving ASD (Pereira et al., 2024, 2019). These results also evidence a significant lateralization of the functional imaging outcomes. As the visual processing of faces usually presents right lateralization, the results reinforce this notion of the right lateralization of facial emotion processing.

### NF training using pSTS as the target and the impact on the imagery task

Contrary to the localizer data, in which patients were asked to passively view an avatar’s facial expressions, during the NF task, they were asked to imagine facial expressions. The final transfer run does not represent a training run since no feedback is provided to the participants, but rather provides the final outcome functional data.

The network commonly engaged during this type of imagery includes the pSTS and other imagery, control, and visual areas, such as the middle frontal cortex, premotor cortex, lingual gyrus, and parahippocampal gyrus (Kim et al., 2007). Previous research has identified several differences between typical development and ASD regarding imagery. (Kana, 2006) found evidence supporting the underconnectivity theory during an imagery task of sentences.

In the NF training, we proposed a direct link between the visual interface and the imagery task (Simoes et al., 2018). Two of the runs were performed without the visual interface as a control to measure the potential transfer of the training. Using these imagery runs, we analyzed differences between session 1 and session 5 of the NF intervention protocol, which showed a decrease in connectivity between the seed region (target of neurofeedback intervention) and the bilateral Frontal Pole, left Supramarginal gyrus, left angular gyrus, bilateral Parahippocampal gyrus, and left Middle Frontal Gyrus. These results suggest differential engagement of frontal cognitive-control areas and the face-perception network. One explanation is the change in strategy throughout the protocol, i.e., participants required less executive, cognitive control, and attentional demands.

### FC-Behavioral analysis

Regarding the correlation between the clinical measures and the connectivity with the target region, the results emphasize that the best outcome was achieved by the participants who increased the connectivity between pSTS and occipital visual areas, and also along the middle temporal gyrus, regions that define the third visual pathway. These results suggest that ATEC decreases (improvements) are associated with a preference for the social visual pathway, even for the imagery process/training.

### Conclusion

In conclusion, our study suggests that an NF clinical treatment may contribute to a shift towards high-level processing within the third visual pathway for the social visual pathway, evidenced by its correlation with changes in behavioral ATEC scores following the intervention. The observed connectivity shifts in association with improvements in adaptive behavior underscore the potential of neurofeedback to modulate neural circuits relevant to social processing in ASD. Future research leveraging network neuroscience approaches with the results of clinical trials is crucial for a more comprehensive and mechanistic understanding of the neurocomputational underpinnings of treatment-related effects in ASD, in particular regarding neurofeedback (De Vico Fallani & Bassett, 2019).

