## Supplementary Materials for "A Neurofeedback therapy of facial expression recognition in Autism shifts connectivity to higher levels within the third visual pathway in relation to clinical improvements"

### Supplementary Methods

#### rt-fNRI-NF paradigm

Data acquisition was performed on a 3T Siemens Magnetom Tim Trio scanner with a 12-channel head coil at the Portuguese Brain Imaging Network. Each scanning session started with a T1-weighted high-resolution magnetization-prepared rapid acquisition gradient echo sequence for co-registration of functional data (176 slices; TE: 3.42 ms; TR: 2530 ms; voxel size 1 mm3 isotropic, FA: 7°; matrix size: 256 × 256).

After the structural sequence, we performed a functional localizer to identify the target region-of-interest (ROI) using an echo planar imaging sequence (160 volumes, TR = 2 s, TE = 30 ms, flip angle = 75°, 32 slices, matrix size 64 × 70, in-plane voxel size = 3 × 3 mm2, slice thickness = 2.5 mm, and gap of 0.5 mm). This run was followed by four functional runs (with similar parameters, except for the number of volumes: 300 volumes), as described below. In this work, we focus on functional analysis in the localizer run and in the last imagery run (also known as the transfer run), a task similar to neurofeedback training but without feedback.

##### Localizer

The localizer was designed to map the face network, and consisted of a block-based design with 40 blocks of 8 seconds and five different conditions: Neutral—static neutral face; to subtract static aspects of face processing; Happy—morphing face from neutral to happy; Sad—morphing face from neutral to sad; Alternate expressions—alternating between sad and happy; and Moving dots—randomly moving dots, to subtract motion processing. The contrast of interest featured the balanced subtraction of conditions Neutral and Moving dots from emotion expression conditions Happy, Sad, and Alternate.

##### Imagery runs

The localizer was followed by four imagery task-based fMRI runs (300 volumes with the same EPI sequence parameters as the localizer run), the first and last without feedback - we here analyzed the last functional run, the transfer run. Each run consisted of 25 blocks (12 regulation blocks featuring imagery of non-neutral expressions alternating with 13 regulation blocks of neutral expression imagery), each block with a duration of 24 seconds. The block sequence included three randomly presented conditions featuring suggested NF strategies: Happy Imagery, Sad Imagery, and Alternate Imagery, interleaved with active Neutral Imagery. Each run starts with a block corresponding to the Neutral Imagery condition.

Explicit strategies were suggested to patients to modulate pSTS BOLD signal activity, for example, by imagining morphing facial expressions of an avatar or a real person, with or without context. Nevertheless, patients were given the freedom to adjust their strategy to maximize feedback modulation. Patients were instructed to use the most successful strategy in the transfer run.

#### ROI-to-ROI pipeline description

Analyses of fMRI data were performed using CONN[1] (RRID:SCR_009550) release 22.a[2] and SPM[3] (RRID:SCR_007037) release 12.12.6.

Preprocessing: Functional and anatomical data were preprocessed using a modular preprocessing pipeline[4], including realignment with correction of susceptibility distortion interactions, slice timing correction, outlier detection, direct segmentation and MNI-space normalization, and smoothing. Functional data were realigned using the SPM realign & unwarp procedure[5], where all scans were coregistered to a reference image (first scan of the first session) using a least squares approach and a 6-parameter (rigid body) transformation[6], and resampled using b-spline interpolation to correct for motion and magnetic susceptibility interactions. Temporal misalignment between different functional data slices was corrected following the SPM slice-timing correction (STC) procedure[7,8], using sinc temporal interpolation to resample each slice BOLD time-series to a common mid-acquisition time. Potential outlier scans were identified using ART[9] as acquisitions with framewise displacement above 0.9 mm or global BOLD signal changes above 5 standard deviations[10,11], and a reference BOLD image was computed for each subject by averaging all scans excluding outliers. Functional and anatomical data were normalized into standard MNI space, segmented into grey matter, white matter, and CSF tissue classes, and resampled to 2 mm isotropic voxels following a direct normalization procedure[11,12] using SPM unified segmentation and normalization algorithm[13,14] with the default IXI-549 tissue probability map template. Finally, functional data were smoothed using spatial convolution with a Gaussian kernel of 8 mm full-width half maximum (FWHM).

Denoising: In addition, functional data were denoised using a standard denoising pipeline[15], including the regression of potential confounding effects characterized by white matter time-series (5 CompCor noise components), CSF time-series (5 CompCor noise components), motion parameters and their first order derivatives (12 factors)[16], outlier scans (below 24 factors)[10], session and task effects and their first order derivatives (10 factors), and linear trends (2 factors) within each functional run, followed by high-pass frequency filtering of the BOLD timeseries[17] above 0.008 Hz. CompCor[18,19] noise components within white matter and CSF were estimated by computing the average BOLD signal and the largest principal components orthogonal to the BOLD average, motion parameters, and outlier scans within each subject's eroded segmentation masks. From the number of noise terms included in this denoising strategy, the effective degrees of freedom of the BOLD signal after denoising were estimated to range from 542.1 to 609.8 (average 592.4) across all subjects[11].

First-level analysis networks_morph: ROI-to-ROI connectivity (RRC) matrices were estimated to characterize the functional connectivity between each pair of regions among 14 ROIs. Functional connectivity strength was represented by Fisher-transformed bivariate correlation coefficients from a general linear model (weighted-GLM[20]), estimated separately for each pair of ROIs, characterizing the association between their BOLD signal timeseries. Individual scans were weighted by a boxcar signal characterizing each individual task or experimental condition convolved with an SPM canonical hemodynamic response function and rectified.

Group-level analyses were performed using a General Linear Model (GLM[21]). For each individual connection a separate GLM was estimated, with first-level connectivity measures at this connection as dependent variables (one independent sample per subject and one measurement per task or experimental condition, if applicable), and groups or other subject-level identifiers as independent variables. Connection-level hypotheses were evaluated using multivariate parametric statistics with random-effects across subjects and sample covariance estimation across multiple measurements. Inferences were performed at the level of individual clusters (groups of similar connections). Cluster-level inferences were based on parametric statistics within- and between- each pair of networks (Functional Network Connectivity[22]), with networks identified using a complete-linkage hierarchical clustering procedure[23] based on ROI-to-ROI anatomical proximity and functional similarity metrics[24]. Results were thresholded using a combination of a $p < 0.05$ connection-level threshold and a familywise corrected p-FDR < 0.05 cluster-level threshold[25].

### Seed-to-Voxel pipeline description

Analyses of fMRI data were performed using CONN[1] (RRID:SCR_009550) release 22.a[2] and SPM[3] (RRID:SCR_007037) release 12.12.6.

Preprocessing: Functional and anatomical data were preprocessed using a modular preprocessing pipeline[4] including realignment with correction of susceptibility distortion interactions, slice timing correction, outlier detection, direct segmentation and MNI-space normalization, and smoothing. Functional data were realigned using SPM realign & unwarp procedure[5], where all scans were coregistered to a reference image (first scan of the first session) using a least squares approach and a 6 parameter (rigid body) transformation[6], and resampled using b-spline interpolation to correct for motion and magnetic susceptibility interactions. Temporal misalignment between different slices of the functional data was corrected following SPM slice-timing correction (STC) procedure[7,8], using sinc temporal interpolation to resample each slice BOLD timeseries to a common mid-acquisition time. Potential outlier scans were identified using ART[9] as acquisitions with framewise displacement above 0.9 mm or global BOLD signal changes above 5 standard deviations[10,11], and a reference BOLD image was computed for each subject by averaging all scans excluding outliers. Functional and anatomical data were normalized into standard MNI space, segmented into grey matter, white matter, and CSF tissue classes, and resampled to 2 mm isotropic voxels following a direct normalization procedure[11,12] using SPM unified segmentation and normalization algorithm[13,14] with the default IXI-549 tissue probability map template. Last, functional data were smoothed using spatial convolution with a Gaussian kernel of 8 mm full width half maximum (FWHM).

Denoising: In addition, functional data were denoised using a standard denoising pipeline[15] including the regression of potential confounding effects characterized by white matter timeseries (5 CompCor noise components), CSF timeseries (5 CompCor noise components), motion parameters and their first order derivatives (12 factors)[16], outlier scans (below 24 factors)[10], session and task effects and their first order derivatives (10 factors), and linear trends (2 factors) within each functional run, followed by high-pass frequency filtering of the BOLD timeseries[17] above 0.008 Hz. CompCor[18,19] noise components within white matter and CSF were estimated by computing the average BOLD signal as well as the largest principal components orthogonal to the BOLD average, motion parameters, and outlier scans within each subject's eroded segmentation masks. From the number of noise terms included in this denoising strategy, the effective degrees of freedom of the BOLD signal after denoising were estimated to range from 542.1 to 609.8 (average 592.4) across all subjects[11].

First-level analysis SBC_target: Seed-based connectivity maps (SBC) were estimated characterizing the spatial pattern of functional connectivity with a seed area. Seed regions included Neurofeedback-Target. Functional connectivity strength was represented by Fisher-transformed bivariate correlation coefficients from a weighted general linear model (weighted-GLM[20]), estimated separately for each seed area and target voxel, modeling the association between their BOLD signal timeseries. Individual scans were weighted by a boxcar signal characterizing each individual task or experimental condition convolved with an SPM canonical hemodynamic response function and rectified.

Group-level analyses were performed using a General Linear Model (GLM[21]). For each individual voxel a separate GLM was estimated, with first-level connectivity measures at this voxel as dependent variables (one independent sample per subject and one measurement per task or experimental condition, if applicable), and groups or other subject-level identifiers as independent variables. Voxel-level hypotheses were evaluated using multivariate parametric statistics with random-effects across subjects and sample covariance estimation across multiple measurements. Inferences were performed at the level of individual clusters (groups of contiguous voxels). Cluster-level inferences were based on parametric statistics from Gaussian Random Field theory[22,23]. Results were thresholded using a combination of a cluster-forming $p < 0.05$ voxel-level threshold, and a familywise corrected p-FDR < 0.05 cluster-size threshold[24].

### Supplementary Results

#### 1. Connectivity analysis of neurofeedback training impact on the face network

##### 1.1. ROI-to-ROI analysis (fig 1)

Network Analysis: Connectivity Results

- Significance: p(FWE) = 0.026

| Connection (ROIs) | Statistic | p-unc | p-FDR |
|---|---|---|---|
| Left FFA — Right mSTS | T(14) = 6.24 | 0.000022 | 0.001966 |
| Left pSTS — Right mSTS | T(14) = 3.26 | 0.005718 | 0.260188 |
| Left pSTS — Left OFA | T(14) = -2.95 | 0.010514 | 0.318913 |
| Right pSTS — Right OFA | T(14) = -2.45 | 0.027974 | 0.509118 |
| Right pSTS — Left OFA | T(14) = -2.29 | 0.037935 | 0.575345 |

#### 1.2. Seed-to-Voxel analysis and parametric changes of connectivity over time

##### 1.2.1. Group seed-based (considering the pSTS as the seed region) connectivity analysis, comparing last > first neurofeedback session (fig. 2)

| Cluster (x, y, z) | Size | size p-FWE | size p-FDR | size p-unc | peak p-FWE | peak p-unc |
|---|---|---|---|---|---|---|
| +08,-84, -26 | 1785 | 0.000000 | 0.000000 | 0.000000 | 0.992034 | 0.000016 |
| -38, -82, -36 | 1360 | 0.000012 | 0.000006 | 0.000000 | 0.988189 | 0.000015 |
| +60, +06, +28 | 477 | 0.085419 | 0.025887 | 0.000153 | 1.000000 | 0.000217 |
| -30, +52, +24 | 461 | 0.103082 | 0.025887 | 0.000187 | 0.999997 | 0.000063 |
| -12, -16, -30 | 392 | 0.230655 | 0.049915 | 0.000450 | 1.000000 | 0.002591 |

##### 1.2.2. Group seed-based (considering the pSTS as the seed region) connectivity analysis, displaying a regression considering the five sessions (fig. 3)

| Cluster (x, y, z) | Size (voxels) | size p-FWE | size p-FDR | size p-unc | peak p-FWE | peak p-unc |
|---|---|---|---|---|---|---|
| +06, -84, -24 | 3635 | 0.000000 | 0.000000 | 0.000000 | 0.882548 | 0.000005 |

#### 1.3. Relation between functional connectivity and clinical outcomes (fig. 4)

| Cluster (x, y, z) | Size | size p-FWE | size p-FDR | size p-unc | peak p-FWE | peak p-unc |
|---|---|---|---|---|---|---|
| -24,-46, -12 | 2413 | 0.000000 | 0.000000 | 0.000000 | 0.999731 | 0.000030 |
| +58, -56, +00 | 2377 | 0.000000 | 0.000000 | 0.000000 | 0.912574 | 0.000006 |

| | | | | | | |
|---|---|---|---|---|---|---|
| -50, -14, -14 | 959 | 0.000434 | 0.000118 | 0.000001 | 1.000000 | 0.000415 |
| -10, +20, -10 | 899 | 0.000788 | 0.000161 | 0.000001 | 1.000000 | 0.000366 |

#### 2. Functional connectivity changes during imagery after neurofeedback training

##### 2.1. Seed-to-Voxel (fig. 5)

| Cluster (x, y, z) | Size | size p-FWE | size p-FDR | size p-unc | peak p-FWE | peak p-unc |
|---|---|---|---|---|---|---|
| -46, +44, +02 | 1147 | 0.000727 | 0.000359 | 0.000002 | 1.000000 | 0.000723 |
| +26, +62, +02 | 1141 | 0.000762 | 0.000359 | 0.000002 | 1.000000 | 0.000548 |
| -44, -72, +44 | 942 | 0.003762 | 0.001183 | 0.000008 | 1.000000 | 0.000530 |
| -06, +08, -22 | 866 | 0.007121 | 0.001682 | 0.000015 | 0.999980 | 0.000070 |
| -36, +34, +40 | 555 | 0.115859 | 0.023179 | 0.000263 | 1.000000 | 0.000552 |

##### 2.2. Relation between functional connectivity and behavioral outcomes (fig. 6)

| Cluster (x, y, z) | Size | size p-FWE | size p-FDR | size p-unc | peak p-FWE | peak p-unc |
|---|---|---|---|---|---|---|
| +24, -76, +18 | 6860 | 0.000000 | 0.000000 | 0.000000 | 0.857491 | 0.000006 |
| +50, -44, +12 | 766 | 0.016070 | 0.006857 | 0.000034 | 1.000000 | 0.000751 |
